# Predicting serious rare adverse reactions of novel chemicals

**DOI:** 10.1101/160473

**Authors:** Aleksandar Poleksic, Lei Xie

## Abstract

Adverse drug reactions (ADRs) are one of the main causes of death and a major financial burden on the world’s economy. Due to the limitations of the animal model, computational prediction of serious, rare ADRs is invaluable. However, current state-of-the-art computational methods do not yield significantly better predictions of rare ADRs than random guessing. We present a novel method, based on the theory of “compressed sensing”, which can accurately predict serious side-effects of candidate and market drugs. Not only is our method able to infer new chemical-ADR associations using existing noisy, biased, and incomplete databases, but our data also demonstrates that the accuracy of our approach in predicting a serious adverse reaction (ADR) for a candidate drug increases with increasing knowledge of other ADRs associated with the drug. In practice, this means that as the candidate drug moves up the different stages of clinical trials, the prediction accuracy of our method will increase accordingly. Thus, the compressed sensing based computational method reported here represents a major advance in predicting severe rare ADRs, and may facilitate reducing the time and cost of drug discovery and development.

---

Adverse drug reactions (ADRs) are one of the main burdens in modern drug discovery [1]. Rare and serious ADRs are responsible for failed drug discovery pipelines and for drug market withdrawals. Cumulative costs of the management of ADRs have been estimated at more than 30 billion per year in the US alone [2]. Clinical impact, including the emergency department visits and prolonged hospital stay, account for a large portion of health care cost. Up to one-third of emergency visits by older adults are due to ADRs [3], while more than one third of ADRs in the pediatric population are potentially life threatening [4]. According to a nationwide Swedish study, ADRs rank 7^th^ among all causes of death [5]. The figures from US studies are even more alarming as they place ADRs as the 4^th^ most common cause of death, ahead of diabetes, pulmonary disease, AIDS, pneumonia, general accidents, and automobile accidents [6].

Finding ADRs for a drug before the drug reaches the market is a difficult and an error prone task. The results of testing a chemical on animals do not always correlate to those obtained when testing the same chemical on humans. Moreover, the patient population recruited during clinical trials is small and biased and hence the data is not statistically robust. Most importantly, clinical trials fail to identify rare and serious side-effects, due to relatively small study duration.

Post marketing surveillance allows for a statistically significant patient population that is followed for a longer period of time. However, the results of post-marketing studies are mostly based on combination drugs and thus are difficult to interpret. More specifically, it is challenging to tell which drug (among multiple ones given to the patient) gives rise to the reported side-effect.

Recent years have seen development of computational approaches to predicting adverse drug reactions. Mizutani et al. [7] employ “canonical correlation analysis” using the information about chemical substructures and drug’s protein targets. Huang et al. [8] use the Support Vector Machines to integrate chemical substructures and protein-protein interaction (PPI) networks. Bresso et al. [9] apply machine learning on integrated functional annotation, pathways and drug characteristics to predict and understand ADR mechanisms. Liu et al. [10] use machine-learning to integrate drugs’ characteristics, such as indications and known ADRs, with the drug's chemical structures, known targets and pathways. Zhang et al. [11] view ADR prediction as a multi-label and ensemble learning task. In their ML algorithm, drug features are associated to side-effects while feature dimensions represent biological components. Xiao et al. [12] apply Symbolic Latent Dirichlet Allocation to learn hidden topics that represents biochemical mechanism that associates drugs to ADRs.

While the advances in the area of computational ADR prediction are encouraging, the field is still at its infancy when it comes to predicting rare and serious ADRs. A harmful ADR often surfaces years or even decades after the drug has been approved. An inability to predict these events leads to complications in diseases and treatments, which can have long term consequences and fatal outcomes. Drug pipeline failures and post-marketing drug withdrawals result in loss of effective compounds (those for which the benefit-to-harm balance is unfavorable), which in turn results in loss of revenue by the drug manufacturer. A methodology capable of predicting ADRs, long before the drug reaches the market or even before the drug is withdrawn from the market, would significantly enhance drug discovery and improve human health.

We show that a variant of the “compressed sensing” technique (namely the “low-rank matrix completion”), from the digital signal processing field, can be easily adapted and used to predict drug-ADR associations with unmatched accuracy. Originally proposed to solve problems arising in coding and data acquisition, compressed sensing (CS) has proved to be an efficient way of recovering any type of signal from few and erroneous samples [13,14,15]. In the framework of ADR prediction, the “signal” can be thought of as the set of all drug-ADR associations (those already known and those yet to be found). The “sample” represents known (reported) associations, identified and stored in the existing drug-ADR databases, such as SIDER [16]. The key observation is that the sample, defined this way, is both sparse and noisy, due to the well-known difficulty of identifying ADRs during clinical trials and post-marketing studies. Therefore, just like the problems in imaging and face recognition, or problems in optical systems research or wireless networking, the drug-ADR association prediction problem is highly amenable to “compressed sensing” solution.

## RESULTS

The compress sensing (CS) algorithm is able to reliably infer new chemical-ADR associations using existing noisy, biased, and incomplete data stored in the SIDER database [16]. Not only is CS highly tolerant to database errors (mislabeled drug-ADR associations), but it also handles sparse data (yet unknown/unrecorded associations) well. Our method is particularly accurate in predicting severe rare ADRs in cases where some (but not necessarily rare) ADRs for the drug are already known.

We compared our approach to two recent state-of-the-art algorithms for drug-ADR association prediction: “multi-label” learning” (ML) [11] and “canonical correlation analysis” (CCA) [7]. In recent benchmarking studies, these two methods exhibited superior accuracy when compared to many other methods for the same problem [11]. For a fair comparison, the benchmarks presented here use the same tests protocol and the same dataset as employed in [11].

In an attempt to gain insight into the progress in the field of computational prediction of serious ADRs, we also submitted to our benchmark a naïve, reference method (here called REF). This straightforward method sets the probability that a given drug will give rise to a particular ADR to the overall promiscuity of that ADR. In other words, the probability of a side-effect *A* being associated to any drug is the same for all drugs and is set to the overall frequency of occurrence of *A*.

### Description of the test set and benchmarking measures

To perform a comprehensive analysis of the methods’ performances, we ran several different cross-validation tests on the set of all drug-ADR associations from the SIDER database. SIDER (version 4.1) contains drug-ADR association data for 1430 FDA approved drugs and 5868 ADRs. Drug-ADR association data from SIDER is represented as a binary matrix *R*, where an entry *r*_*i,j*_ is equal to 1 if the drug *i* is known to cause ADR *j* and 0 otherwise.

Each method submitted to our benchmarks was run using its default parameters. To make ADR predictions, some methods (namely CS and ML) take advantage of the pairwise similarity of drugs in the test set. Consistent with the procedure in [11], we provided those methods with the same matrix of Tanimoto similarity (Jaccard index) scores [17] between pairs of drugs. In contrast to CS and ML, the CCA algorithm takes encoding of drugs’ chemical structures as input. To ensure a fair comparison, we supplied CCA with the same set of chemical fingerprints used by the authors of the CCA algorithm, namely the set of 881 chemical substructures specified in the PubChem database [18].

Our tests employ two classical performance measures, namely the area under the ROC curve (AUC) and the area under the PR curve (AUPR). The ROC (Receiver Operating Characteristic) curve represents the relationship between the false-positive and the true-positive rate while the PR (Precision-Recall) curve represents the relationship between the sensitivity (true positive rate or recall) and the positive predictive value (precision).

### Assessing current state of the field of computational ADR prediction

To assess methods’ accuracies on rare ADRs, we first ran multiple, independent and statistically robust, cross-validation experiments, one for each selected ADR promiscuity cutoff (12, 25, 50, 100, 200, 400, 800, ∞), where the “promiscuity” of an ADR is defined as the number of FDA approved drugs that are known to cause the ADR. For each promiscuity cutoff *x*, a cross-validation experiment was performed on the set of drug-ADR pairs in which the ADR’s promiscuity does not exceed *x*.

Our analysis presented in Fig. 1 not only confirms the published accuracy of current methods but also provides an insight into the performance of the state-of-the-art algorithms as well as the accuracy improvement offered by compressed sensing (CS). We can summarize the results shown in Fig. 1 as follows:

i. <u>The existing algorithms are unable to predict serious rare side-effects</u>. While published accuracies of current methods are more or less satisfactory (balanced and unbalanced AUC and balanced AUPR ~0.9; unbalanced AUPR ~0.35), they should be interpreted properly, since they only represent the average accuracies computed for all ADRs combined (the right side of Fig. 1). The cumulative accuracies are driven strongly by easy predictions of frequent and innocuous ADRs, those of little interest in drug discovery. For rare and serious ADRs, the accuracy of current algorithms quickly approaches the accuracy a purely random classifier (AUC ~0.5).
ii. <u>To date, the progress in the field of computational ADR prediction has been dismal at best</u>. To assert this claim, it is enough to glance over the line that traces the performance of the naïve and straightforward REF method in Fig 1. Going beyond this simple approach and implementing more sophisticated techniques, such as multi-label learning and canonical correlation analysis, yields a low diminishing return.
iii. <u>“Compressed sensing” (CS) overcomes current obstacles in predicting drug-ADR associations</u>. Our method is so efficient in extracting relevant information from noisy, biased and incomplete data (stored in the SIDER database) that its performance in predicting severe ADR (left part of Fig. 1) matches or even exceeds the cumulative performances of current methods on all side-effects combined (right part of Fig. 1).

**Fig. 1.**
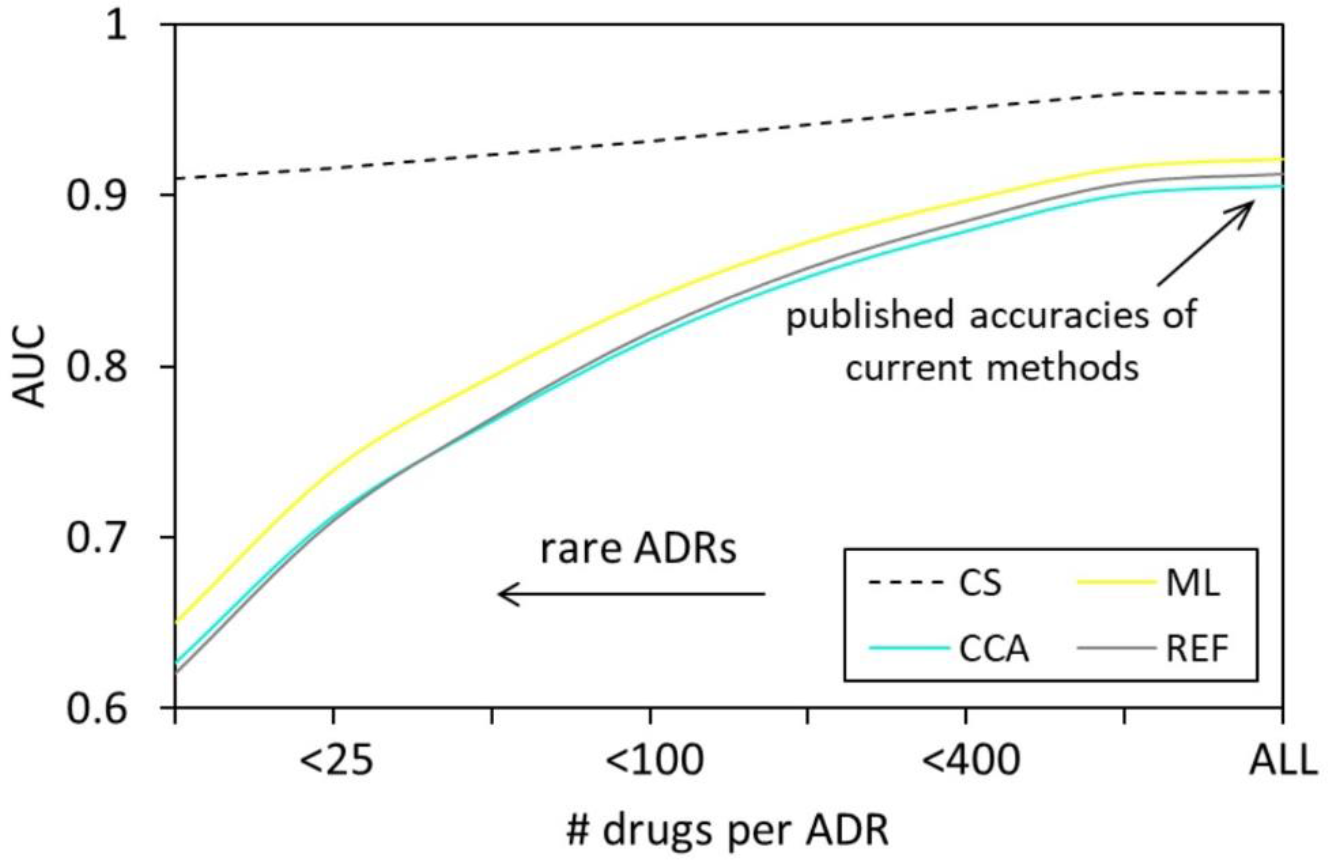
State-of-the-art in ADR prediction and the value added by compressed sensing. CS: Compressed Sensing; ML: Multi-Labeling; CCA: Canonical Correlation Analysis; REF: naïve (reference) method. The values on the x-axis represent ADR promiscuities. The *y*-axis represents the performance metrics, defined as the area under the ROC curve (AUC). The results were obtained using a statistically rigorous cross-validation experiment (STDERR too small to show).

### Compressed sensing learns on the fly

Not only is compressed sensing able to predict rare ADRs, but also (as demonstrated later) the performance of CS in predicting ADRs for a particular chemical improves with the increasing knowledge of other ADRs associated with the chemical. In practice, this means that the ability of compressed sensing to predict a serious ADR for a candidate chemical would increase as the drug moves up the different stages of clinical trials. Other methods are unable to take advantage of this information. This comes as no surprise to us, since a closer look into the ML algorithm reveals that, when predicting whether a drug *D* is likely to cause an ADR *A*, ML utilizes the information on other drugs that cause the side-effect *A*, but not the information on other ADRs associated to *D*.

Before running a more comprehensive benchmark, we tested the performance of compressed sensing (CS) in predicting selected serious side-effects, including hepatotoxicity, cardiotoxicity, carcinogenicity, neurotoxicity, as well as thrombocytopenia, leukopenia, anaemia, nepropathy and death. Those nine ADRs represent some of the main causes of drug market withdrawals [19]. Starting with hepatotoxicity, we selected all drugs known to cause this ADR (“cases”) and the same number of randomly selected drugs that are known not to cause hepatotoxicity (“controls”). Fig. 2 illustrates the differences in normalized raw scores obtained by the compressed sensing (CS), ML and CCA on “case” and “control” drugs.

The performance statistics (AUC and AUPR) obtained from the algorithms’ raw scores (after averaging the raw scores over a dozen of randomly chosen sets of “control” drugs) is presented in Fig. 3. Summary performance data for cardiotoxicity, hepatotoxicity, and neurotoxicity is presented in Fig. 4. As shown in the Supplementary figure S1, the results for the remaining six ADRs show similar trends.

**Fig. 2.**
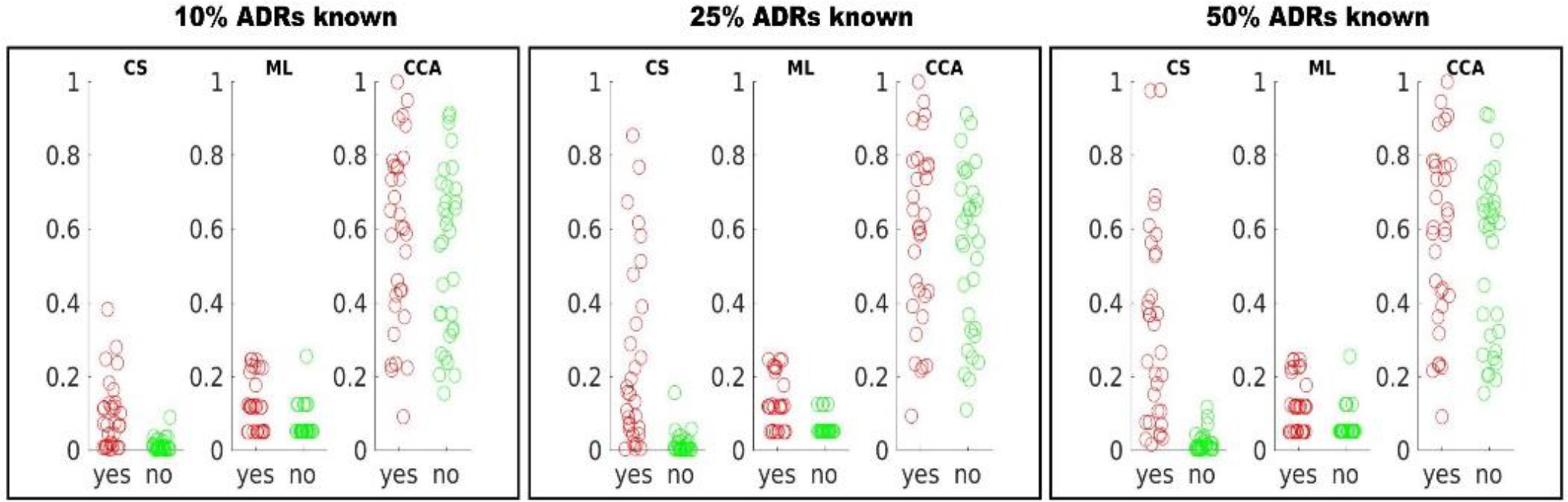
Predicting hepatotoxicity of drugs. Drugs known to cause hepatotoxicity (“cases”) are shown as red circles. Green circles (“controls”) represent randomly chosen drugs known not to cause hepatotoxicity. The vertical axis gives normalized prediction scores. It is obvious that only compressed sensing (CS) can differentiate between the two sets of drugs. It should be noted that the method’s performance might be better than suggested in the figures. For instance, the “control” outlier (green circle in the middle figure), corresponding to the drug *minoxidil*, clearly stands out by its high CS score. Despite being classified as a non-hepatotoxicity drug in SIDER, *minoxidil* (according to FDA case reports), in fact, is known to cause hepatotoxicity in some patients, particularly those over the age of 60.

**Fig. 3.**
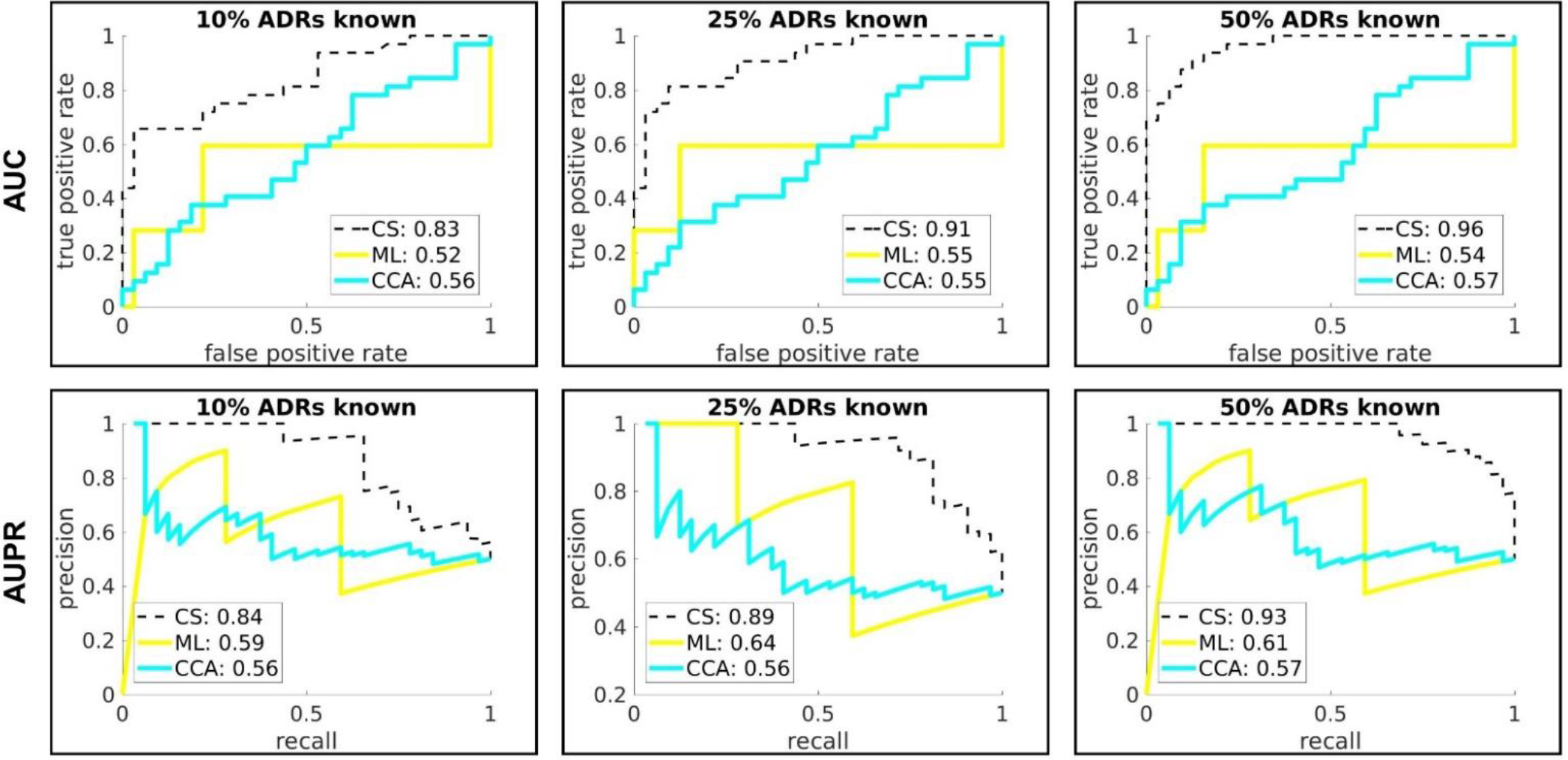
Accuracy of hepatotoxicity predictions. The ROC (top) and PR (bottom) curves are generated based upon the raw scores obtained on “case” and “control” drugs. We performed a number of different tests, each time letting the methods under study (CS, ML, CCA) access different amount of information (10%, 25%, 50%) on other, non-hepatotoxicity ADRs associated to “case” and “control” drugs, thus mimicking methods’ accuracy and reliability during clinical trials. We use “balanced” AUPR for better visualization. Unbalanced AUPR scores are easily obtained by multiplying the balanced scores by the fraction of condition positives in the test set.

**Fig. 4.**
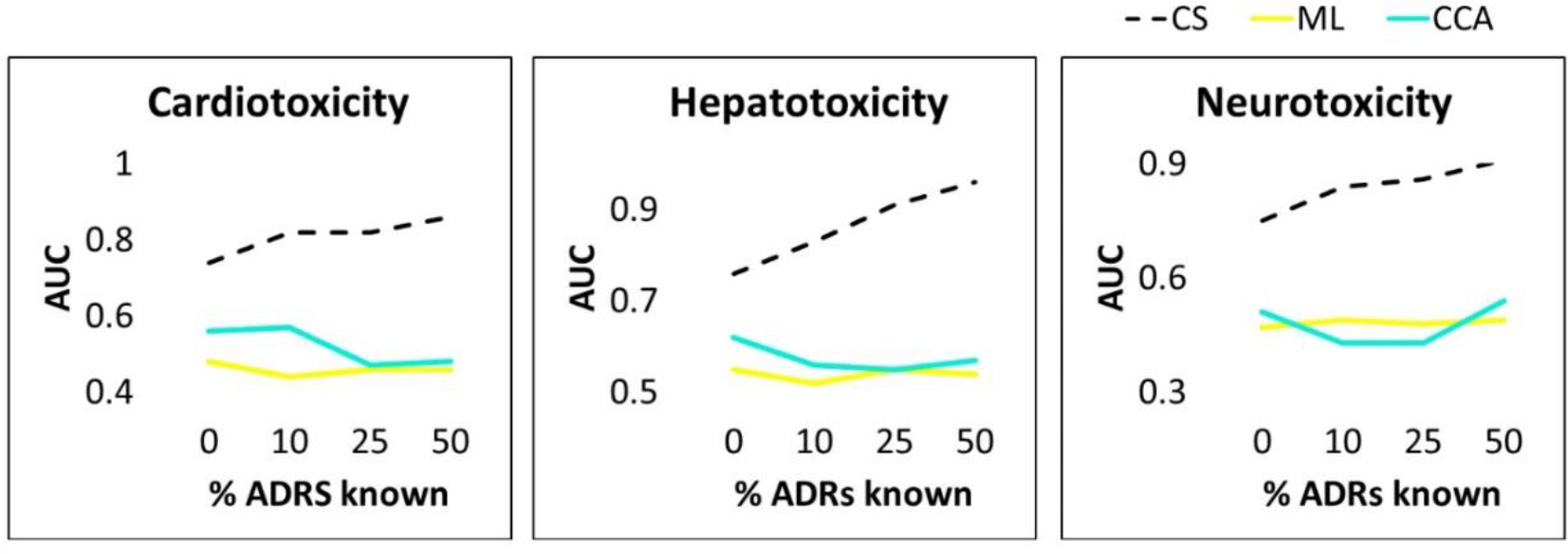
Predicting ADRs responsible for drug market withdrawals. The x-axis represents the percentage (0%, 10%, 25%, 50%) of already known ADRs for the drug. The y-axis represents the AUC values. The mean AUC values shown in the figures are obtained over multiple runs on “control” drugs to achieve robust statistics (STDERR too small to show). Corresponding figures for other selected ADRs are given in the supplementary material.

### Comprehensive cross-validation benchmarks

We now return to the comprehensive benchmark from the beginning of this section to provide a more detailed and more illustrative performance analysis, illustrated in Fig. 5 and Fig. 6. For better performance visualization, figures 5 and 6 show a head-to-head comparison between the maximum possible accuracy scores obtained by current, State-Of-the-Art (SOA) methods and corresponding scores obtained by CS.

**Fig. 5.**
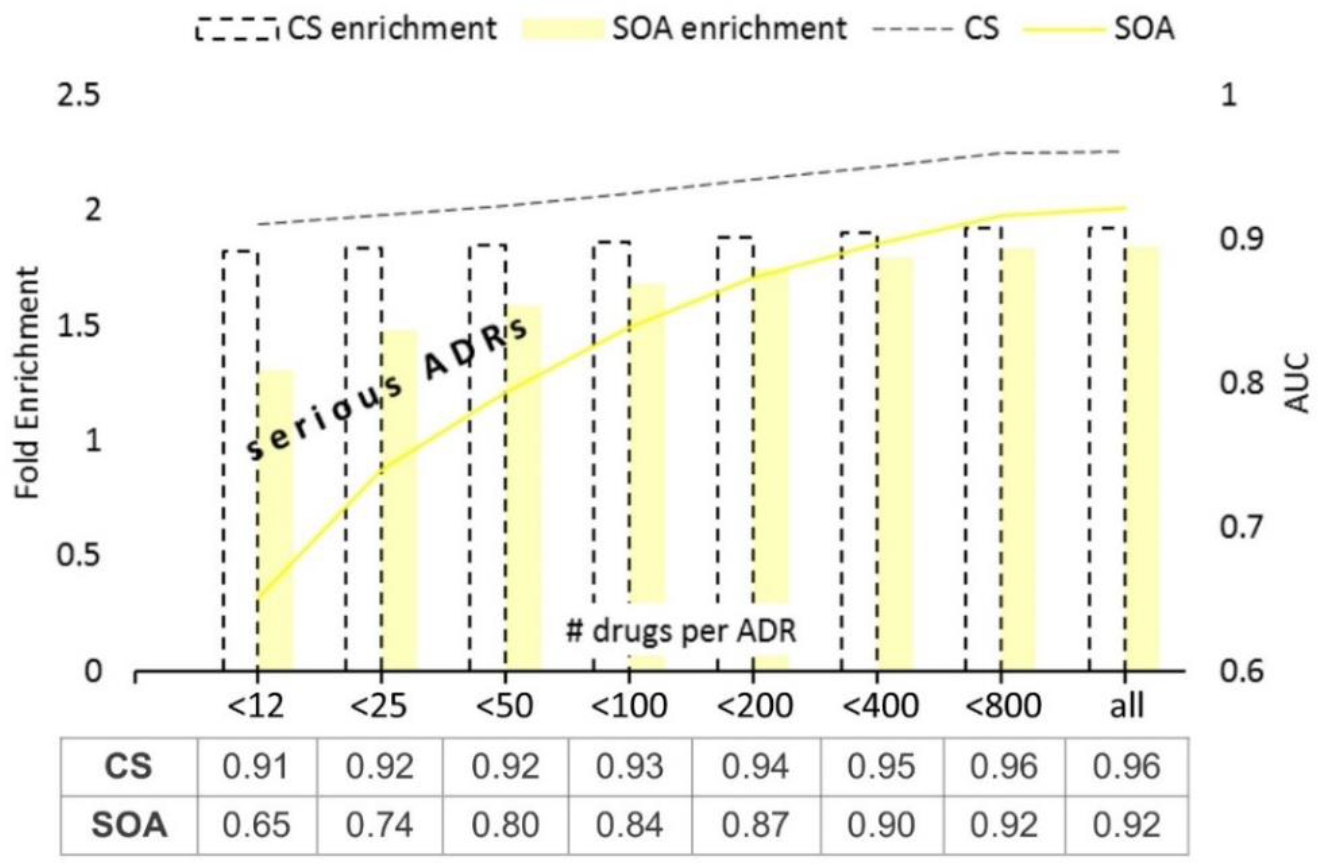
Value added by compressed sensing (CS) in the AUC benchmark. For clarity of presentation, we only show the comparison between CS scores and SOA scores, where the SOA (State-Of-the-Art) score in each test category (<12, <25, etc.) is defined as the maximal AUC score among all AUC scores obtained by state-of-the-art methods in that particular category. The data table beneath the graph gives the actual AUC values for CS and SOA. STDERR is too small to show.

Aside from showing the raw scores, Fig. 5 and Fig. 6 illustrate the “fold enrichment” offered by CS and SOA. The “fold enrichment” represents the improvement in a method’s performance over the random predictor (one that generates prediction scores uniformly at random). In other words, the “fold enrichment” shows how many times is the method’s AUC (or AUPR) better than the AUC (respectively, AUPR) obtained by the purely random classifier. Fold enrichment is particularly useful when interpreting the AUPR scores (Fig. 6), because, in contrast to the intuitive AUC scores, the AUPR scores depend on the property of the test set. More precisely, the AUPR scores obtained from the purely random classifier are equal to the fraction of condition positives in the test set.

**Fig. 6.**
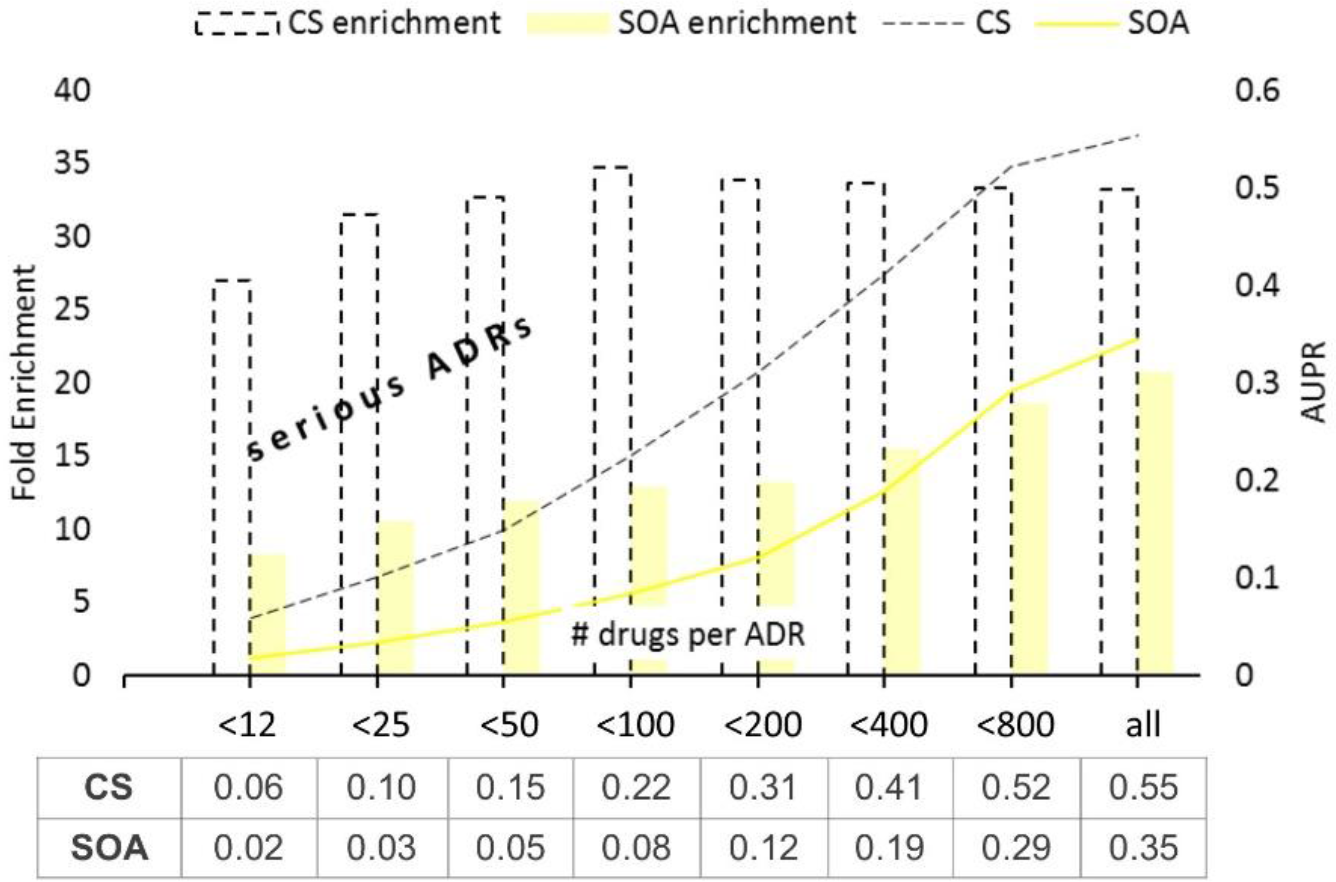
Value added by compressed sensing (CS) in the AUPR benchmark. STDERR is too small to show. In contrast to AUC, the actual AUPR values (shown in the data table beneath the graph) are more difficult to interpret. To put them in the context, we note that the AUPR of a random classifier represents the fraction of conditional positives in the data set (∑ *cond.pos* / (∑ *cond.pos* + ∑ *cond.neg*)). We also note that the performance of both CS and SOA in this benchmark is probably much better than shown, due to unreliability and sparseness of data in the SIDER database (we refer the reader again to the illustrative example provided in the caption of Fig. 2).

As seen in Fig. 5 and Fig. 6, CS enriches prediction of ADRs at an almost uniform rate, irrespective of the ADR promiscuity and the type of test performed (AUC or AUPR). For extremely rare ADRs, those associated with less than 12 FDA drugs (such as *carcinogenicity* or *death*), the performance of CS, as measured by AUPR, is about 27 times better than the performance of the random classifier, while the performance of SOA is only about 8 times better. For the more frequent serious ADRs (associated with < 50 drugs), such as *neurotoxicity* or *cardiotoxicity*, the fold enrichments of CS and SOA are 34 and 12, respectively.

### Predicting ADRs for chemicals with a few or no known rare ADRs

The performance of compressed sensing on ADRs responsible for drug market withdrawals, presented earlier in Fig. 4, suggests that CS might have significant advantage over the other methods, even in cases of chemicals with no known ADRs (left side of the Fig. 4 plots). Furthermore, the same figure suggests that such advantage will increase with the increasing number of ADRs discovered for the drug (right side of the Fig. 4 plots). To test this hypothesis on a large scale, we ran a comprehensive cross-validation benchmark. Specifically, for each ADR promiscuity cutoff, we removed (and then tried to re-discover) all associations between drugs and their severe, rare ADRs (those with promiscuity below the cutoff). The results of our analysis are given in Fig. 7. As seen in this figure, the AUPR fold enrichment achieved by CS is about three times as large as the one achieved by SOA.

**Fig. 7.**
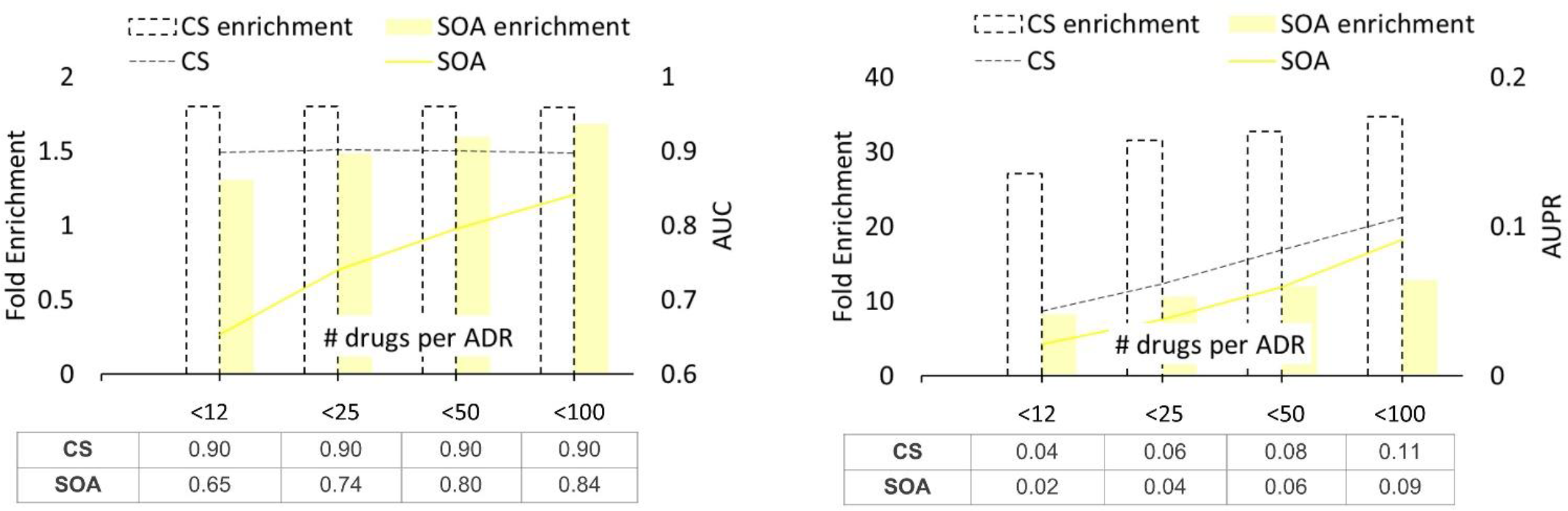
Value added by compressed sensing (CS) in the AUC (left) and AUPR (right) test for rare ADRs. The actual AUC and AUPR scores represent the mean values obtained from 5 rounds of the 10-fold cross-validation test on the set of “new” drugs; those with all rare ADRs masked out (hidden). STDERR values are too small to show.

### Predicting ADRs for chemicals of novel structure

Finally, we used cross-validation to test the ability of our method to correctly predict side-effects of novel chemicals. For the purpose of this study, a chemical is considered to have a “novel” structure if its Tanimoto similarity to every other database chemical is below the upfront specified cutoff.

To perform the cross-validation test, the training set has to be altered by removing all chemicals (along with their ADR associations) that have above the threshold Tanimoto similarity to any chemical from the test set. This intervention renders 10-fold cross validation unfeasible, due to the training set in each fold being nearly (or completely) empty. Hence, in order to gain insight into methods’ performance in discovering rare ADRs for novel drugs, we resorted to leave-one-out-cross-validation (LOOCV) on the set of drugs. For each drug in the test set, we hide, and then try to recover, all rare ADRs (those of drug-promiscuity below the specified cutoff) associated with that drug.

The example Fig. 8 (A) shows a head-to-head comparison between CS and SOA in recovering hidden ADRs associated to less than 25 drugs, using varying Tanimoto cutoffs for excluding “similar” drugs from the training set. As seen in this figure, even if provided a severely reduced training set (Tanimoto cutoff = 0.1) our algorithm exhibits accuracy superior to the accuracies of other state-of-the-art methods even if those methods are trained on comprehensive sets (Tanimoto cutoff = 1.0). Additional data is presented in the Table (B). Complete results of the LOOCV benchmark are given in the supplementary table S1.

**Fig. 8.**
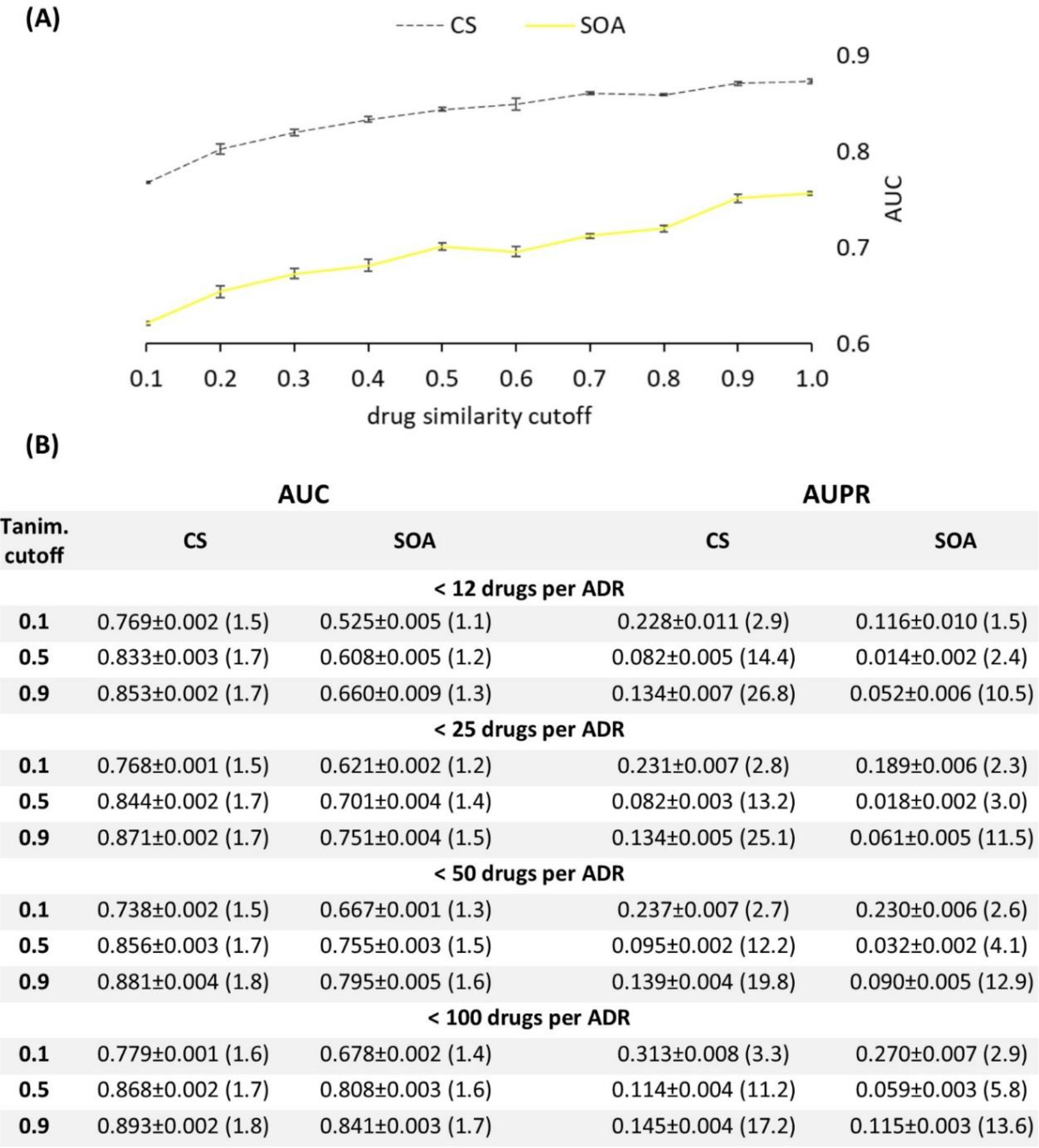
The results of LOOCV for chemicals of novel 3D structure. The actual AUC and AUPR scores represent the mean values obtained from 5 rounds of the leave-one-out-cross-validation test on the sets of 100 randomly chosen drugs. For each drug tested, we removed from the training set all drugs form the same structural class i.e., those that have the above cutoff Tanimoto similarity to the test drug. The fold enrichment over the random classifier is given in parentheses. The top figure (A) shows the AUC values obtained on ADRs associated with <25 FDA approved drugs. The table beneath the figure gives selected methods’ accuracy values for different Tanimoto and ADR promiscuity cutoff values. Complete table can be found in the supplementary material.

## METHODS

Starting from a known (in practice, noisy and incomplete) binary matrix of drug-ADR associations *R* = (*r*_*i,j*_), the pairwise ADR similarity matrix *M* = (*m*_*i,j*_) and the pairwise drug similarity matrix *N* = (*n*_*i,j*_), our algorithm outputs the “latent” ADR and drug preferences *F* = (*f*_*i,j*_) and *G* = (*g*_*i,j*_) by minimizing the loss function

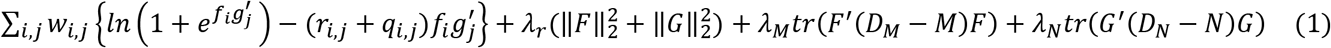

In the function (1) above, *f*_*i*_ represents the *i*^*th*^ row of *F, F′* is the transpose of *F* and ∥ ∥_2_ represents the Frobenius norm. We use *tr* to denote the “matrix trace” and *D*_*M*_ to denote the “degree matrix” of *M* (namely, the diagonal matrix whose diagonal element in row *i* represents the sum of all elements of *M* that belong to row *i*). The lambdas (*λ*) are optimizable parameters. The output matrix of drug-ADR associations is computed as *P* = exp(*FG*′)/(1 + exp(*FG*′)), where exp() represents the matrix exponential. A schematic diagram illustrating the flow of our algorithm is given in Fig. 9.

**Fig. 9.**
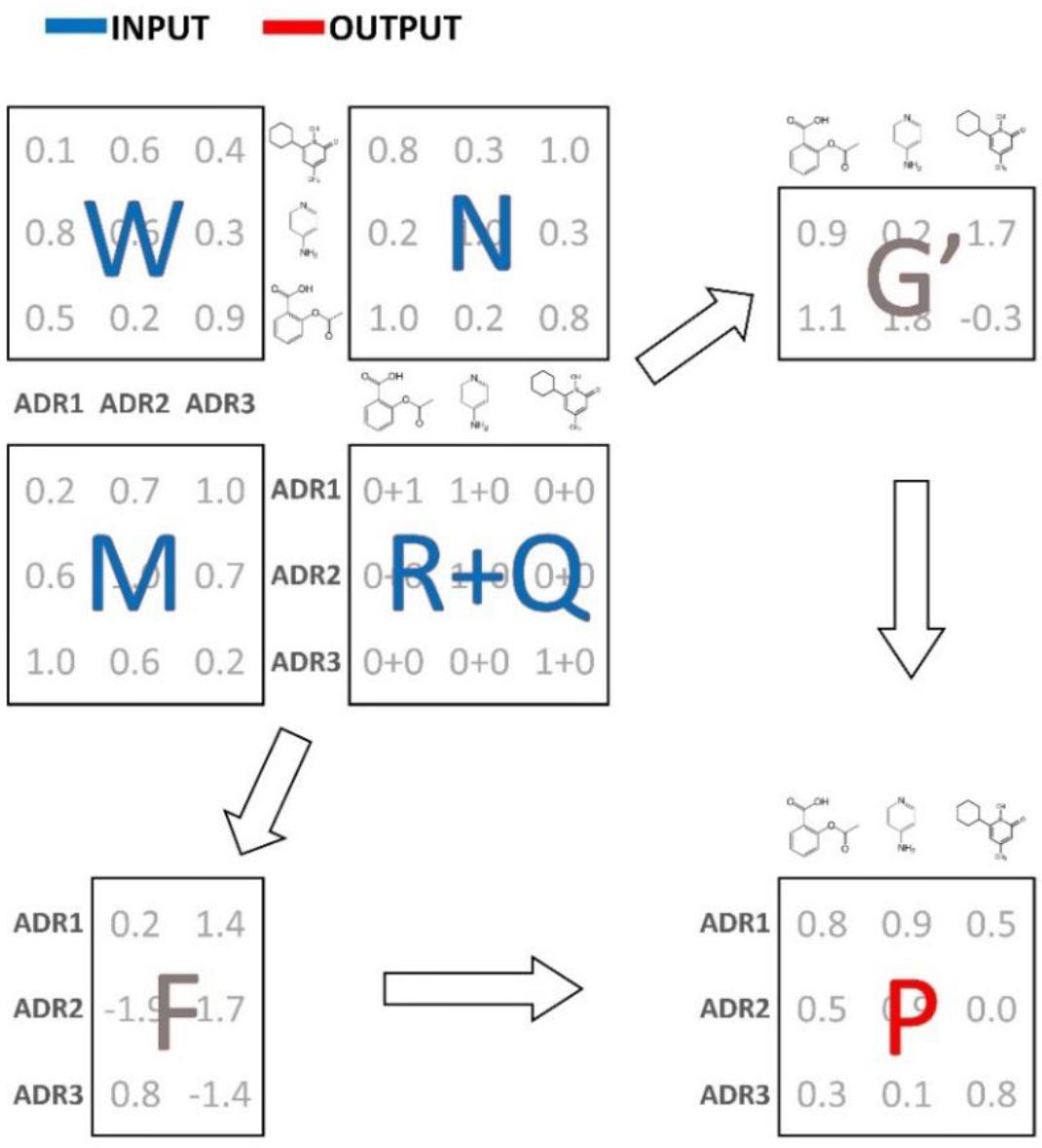
Algorithm flow. R: known drug-ADR associations (sample); M: pairwise ADR similarity matrix; N: pairwise drug similarity matrix; W: drug-ADR frequencies; Q: impute values; F: latent ADR preferences; G: latent drug preferences; P: output drug-ADR probabilities (recovered signal). Algorithm’s input and output are shown in blue and red colors, respectively.

The first two terms in (1) drive the “signal recovery” process (matrix completion), whereas the last two terms mandate that similar drugs have similar side-effects and vice versa. Although our method is capable of factoring in the drug-ADR frequency values *w*_*i,j*_ and the drug-ADR impute values *q*_*i,j*_ this information is currently not been taken advantage of and *w*_*i,j*_ and *q*_*i,j*_ are set to 1’s and 0’s, respectively.

The matrices of latent ADR and drug preferences (*F* and *G*, respectively) are found during the standard minimization procedure. For the sake of brevity, we skip technical details (see [20]), but emphasize that the key idea behind our approach is to demand that *F* and *G* are small in one dimension. That way, the output matrix *P* of predicted interaction probabilities (recovered signal) must be of small rank and, in turn, free of noise. An efficient optimization of the objective function (1) is achieved using a stochastic gradient descent method [21].

While the pairwise drug similarity scores (*N*) are computed using the classical Jaccard index [22], the notion of pairwise ADR similarity scores (along with the notion of frequencies and impute values) is unique to our method and improves the prediction accuracy. Our pairwise ADR similarity scores are defined as the average of semantic and relatedness measures (*path* and *lesk*, respectively) and are computed by running the *umls-similarity* software [23] on MedDRA vocabulary [24].

An added benefit of our loss function (1) is that it allows one to take advantage of the frequencies of known drug-ADR associations and the drug-ADR impute values. Each *w*_*i,j*_ in the loss function (1) represents the frequency at which the drug *i* causes the side-effect *j*, while each *q*_*i,j*_ can be used to explicitly specify the likelihood of a drug-ADR association. To explain how the impute values can be useful in predicting drug-ADR associations, consider, for instance, an ambiguous case of a newly discovered drug-ADR association that has not yet been recorded in the database (*r*_*i,j*_ = 0). This new knowledge can be easily incorporated into our method by setting *q*_*i,j*_ = 1, while adjusting the corresponding weight *w*_*i,j*_ to account for any uncertainty in the imputed value. However, method uses neither the weight nor the impute value functionality, due to unavailability of comprehensive drug-ADR frequency information in the SIDER database. This might change in the future, as more comprehensive databases, containing frequency information, become available.

While we have originally developed and published the analytical framework (1) for the drug-target interaction problem [20], we subsequently noticed that the compressed sensing is much more amenable to predicting adverse drug reactions (ADRs). In contrast to drug-target interaction problem, where the baseline data is already clean but incomplete, the drug-ADR association data is both, incomplete and noisy. Compressed sensing is particularly suited to deal with such data.

Finally, we note that the running time of our method is comparable to that of CCA but worse than the running time of ML. As is the case with the other algorithms in our study, straightforward parallel implementation can make compressed sensing practical, even in large scale studies.

## DISCUSSION

Adverse drug reactions (ADRs) play a major role in drug discovery and human health. Despite significant efforts made over the last decade, the progress in developing computational tools capable of predicting serious side-effects of novel chemicals and market drugs has been dismal at best. No current computational method is able to predict whether a novel and promising compound will eventually cause hepatotoxicity, carcinogenicity, cardiotoxicity, neurotoxicity, immune reaction thrombocytopenia, leukopenia, anaemia or any other harmful and potentially fatal ADRs. Moreover, advances in the area of drug-ADR association prediction are hindered by a lack of clean and comprehensive databases that store drug-ADR associations and by the difficulty of current methods to deal with noisy and sparse information.

Using the “compressed sensing” framework from the digital signal processing field we developed a computational method that can reliably infer new chemical-ADR associations using existing noisy, biased, and incomplete databases. Originally proposed to address problems in electrical engineering, nowadays “compressed sensing” is a reliable way of recovering any type of signal from a relatively small number of (often erroneous) samples. In the setting of drug-ADR association prediction, the signal can be thought of as the set of all true (known and yet to be discovered) drug-ADR associations. The sample represents known drug-ADR associations, those stored in existing drug-ADR association databases. Because the current databases of drug-ADR associations are sparse and noisy, the drug-ADR association prediction problem is highly amenable to the compressed sensing solution.

Not only is our method able to detect rare ADRs associated with novel chemicals, but also our data demonstrates that the accuracy of compressed sensing in predicting a serious ADR for a candidate drug increases with increasing knowledge of other ADRs associated with the drug. In practice, this means that, as the candidate drug moves up the different stages of clinical trials, the prediction accuracy of our method will increase accordingly.

Compressed sensing represents an important first step in the development of a fully automated and accurate computational method for predicting serious Adverse Drug Reactions (ADRs). Ultimately, accurate and reliable prediction of ADRs will accelerate drug discovery and reduce the risks of drug treatment.

The difficulty in identifying ADRs during clinical trials and the complexity of parsing side-effect data from drug package inserts and post-marketing reports results in creation of incomplete and noisy databases of drug-ADR associations. Cleaner and more comprehensive databases represent a straightforward way of improving the performance of prediction methods. For instance, we were able to increase the accuracy of our method in predicting drug-induced liver injury by replacing SIDER hepatotoxicity associations by those stored in LTKB-BD [25]. While SIDER is an automatically generated and comprehensive source, containing almost 140K drug-ADR associations, LTKB-BD represents an expert classification of only 287 drugs with respect to drug-induced liver injury. Even a quick comparison of the two sets of drugs (comparing only drugs that have easily identifiable Pubchem CIDs) reveals significant classification differences between two sources. For instance, 38 out of 137 “most hepatotoxicity” drugs as well as 56 out of 85 “less hepatotoxicity” drugs in LTKB-BD were labeled as “no known hepatotoxicity” in SIDER.

We also believe that much more accurate predictions of drug-ADR associations can be made by utilizing not only cleaner data on drug-ADR associations but also gender-, age- and demographics-specific drug-ADR associations, drug-dose specific associations, and data on side-effects arising from combination drugs [26].

